# Germ-free mice exhibit mast cells with impaired functionality and gut homing and do not develop food allergy

**DOI:** 10.1101/394213

**Authors:** Martin Schwarzer, Petra Hermanova, Dagmar Srutkova, Jaroslav Golias, Tomas Hudcovic, Marek Sinkora, Johnnie Akgün, Christian Zwicker, Ursula Wiedermann, Ludmila Tuckova, Hana Kozakova, Irma Schabussova

## Abstract

**Background:** Mucosal mast cells (MC) are key players in IgE-mediated food allergy (FA). The evidence on the interaction between gut microbiota, MC and susceptibility to FA is contradictory.

**Objective:** We tested the hypothesis that commensal bacteria are essential for MC migration to the gut and their maturation impacting the susceptibility to FA.

**Methods:** The development and severity of FA symptoms was studied in sensitized germ-free (GF), conventional (CV) and mice mono-colonized with *L. plantarum* WCFS1 or co-housed with CV mice. MC were phenotypically and functionally characterized.

**Results:** Systemic sensitization and oral challenge of GF mice with ovalbumin led to increased levels of specific IgE in serum compared to CV mice. Remarkably, despite the high levels of sensitization, GF mice did not develop diarrhea or anaphylactic hypothermia, common symptoms of FA. In the gut, GF mice expressed low levels of the MC tissue-homing markers CXCL1 and CXCL2 and harbored fewer MC which exhibited lower levels of MC protease-1 after challenge. Additionally, MC in GF mice were less mature as confirmed by flow-cytometry and reduced edema formation after injection of degranulation-provoking compound 48/80. Co-housing of GF mice with CV mice fully restored their susceptibility to develop FA. However, this did not occur when GF mice were mono-colonized with *L. plantarum*.

**Conclusion:** Our results demonstrate that microbiota-induced maturation and gut-homing of MC is a critical step for the development of symptoms of experimental FA. This new mechanistic insight into microbiota-MC-FA axis can be exploited in the prevention and treatment of FA in humans.

## INTRODUCTION

Food allergy (FA) is a widespread pathological immune reaction which is initiated by generally harmless food antigens. Its global prevalence has been increasing since the 1960s, especially in industrialized countries, suggesting environmental factors play a key role in the susceptibility and etiology of this disorder (Yu, Freeland et al. 2016). IgE-mediated FA, which is the most common form of FA, is based on two phases: i) allergic sensitization and ii) the effector phase (Lee, Chen et al. 2016, Yu, Freeland et al. 2016). The humoral and cellular immune responses are clearly biased toward a type 2-related phenotype, characterized by the production of specific IgE antibodies and cytokines, such as IL-4, IL-5, IL-13, or IL-10 (Berin and Sampson 2013). In the effector phase, allergen-induced crosslinking of IgE bound to mast cells (MC) leads to release of histamine, serotonin and MC proteases as well as cytokines, (e.g. TNF-α) resulting in the rapid appearance of symptoms, such as diarrhea and hypothermia (Kraneveld, Sagar et al. 2012). Even in individuals with high levels of food allergen-specific serum IgE, the susceptibility of developing these clinical symptoms differs dramatically (Liu, Jaramillo et al. 2010, Brough, Cousins et al. 2014). The mechanisms underlying this phenomenon remain unknown.

The hygiene hypothesis suggests that changes in the commensal microbiota composition and/or function, due to excessive antibiotic use or increased hygiene, can increase the level of allergic sensitization (Okada, Kuhn et al. 2010, Liu 2015). By using germ-free (GF) animals, several studies have shown that the lack of bacteria leads to increased levels of serum IgE in comparison to colonized conventional (CV) mice (Morin, Fischer et al. 2012, Stefka, Feehley et al. 2014). Interestingly, colonization of GF mice with a single bacterial strain, with a mixture of several strains, or with the microbiota of CV mice through co-housing prevented the development of allergic sensitization and led to decrease of allergen-specific IgE (Schwarzer, Srutkova et al. 2013, Stefka, Feehley et al. 2014, Kozakova, Schwarzer et al. 2016).

*Lactobacillus plantarum* is an extremely versatile lactic acid bacterium that has been isolated from a variety of habitats, such as plants, the gastro-intestinal tracts of human and animals, as well as raw or fermented dairy products (Martino, Bayjanov et al. 2016). The human isolate *L. plantarum* WCFS1 possesses strong immunomodulatory properties, and has been shown to induce maturation of immune cells *in vitro* (Meijerink, Wells et al. 2012, Gorska, Schwarzer et al. 2014) and interact with the host immune system *in vivo* (van den Nieuwboer, van Hemert et al. 2016). Specifically, oral application of *L. plantarum* WCFS1 enhanced activation of intestinal cells and shifted the Th1/Th2 balance towards a Th2 response (Smelt, de Haan et al. 2012). In a mouse model of peanut allergy, oral supplementation of this strain aggravated the allergic responses associated with increased MC degranulation (Meijerink, Wells et al. 2012).

MC are innate immune cells which are involved both in the immunological homeostasis as well as in parasitic infection (Lantz, Boesiger et al. 1998, Pennock and Grencis 2004, Olivera, Beaven et al. 2018) and various immunological disorders (Kurashima and Kiyono 2014, Reber, Sibilano et al. 2015). MC originate from CD34+ progenitors in the bone marrow and then enter the circulation and peripheral tissues, where they undergo maturation (Kunii, Takahashi et al. 2011, Halova, Draberova et al. 2012). Being at the mucosal sites, MC are in close contact with the microbiota. Indeed, commensal bacteria have been shown to modulate several phenotypic and functional characteristics of MC, including their recruitment to the tissue, maturation and survival (Kunii, Takahashi et al. 2011, Wang, Mascarenhas et al. 2016). Along these lines, Kunii *et al*. have shown that the microbiota is required for the migration of MC to the intestine through the induction of CXCR2 ligands (Kunii, Takahashi et al. 2011). Similarly, in the skin, the microbiota is crucial for recruitment and maturation of dermal MC (Wang, Mascarenhas et al. 2016).

Although only low numbers of MC are found in the intestine of naïve mice (Guy-Grand, Dy et al. 1984), their numbers increase in food allergy (Brandt, Strait et al. 2003). The crucial role of MC in FA has been well established (Brandt, Strait et al. 2003, Chen, Lee et al. 2015). After MC depletion with anti-c-kit antibody, CV mice do not develop OVA-induced gastrointestinal manifestation (Brandt, Strait et al. 2003) and MC are also essential for the full development of hypothermia in the OVA FA mouse model (Balbino, Sibilano et al. 2015). Additionally, transgenic mice with increased numbers of intestinal MC exhibit augmented severity of FA symptoms (Ahrens, Osterfeld et al. 2012).

The literature on the interaction between microbiota, MC and susceptibility to FA is contradictory. On one hand, it has been demonstrated that GF mice exhibit altered functionality of MC and their impaired migration into the intestinal and skin tissue (Kunii, Takahashi et al. 2011, Wang, Mascarenhas et al. 2016). On the other hand; different studies have shown that GF mice are more susceptible to develop clinical symptoms of FA (Cahenzli, Köller et al. 2013, Stefka, Feehley et al. 2014).

In this study we seek to determine the role of commensal bacteria in the induction of FA using GF mice. We observed that GF mice did not develop the clinical symptoms of FA, such as allergic diarrhea and hypothermia, despite having higher titers of allergen-specific Th2-associated antibodies. Furthermore, the lack of commensals resulted in reduced numbers of tissue MC with low maturation status. Importantly, conventionalization of GF mice with complex microbiota through co-housing with CV mice, but not mono-colonization with *L. plantarum* WCFS1, fully recapitulated the FA phenotype observed in the CV mice. These results implicate that signals from complex microbiota are necessary for the homing of MC into the intestinal tissue as well as their maturation, which are prerequisites for developing the clinical symptoms of FA.

## METHODS

### Animals

Germ-free (GF) BALB/c mice were derived from the conventional BALB/c mice by Caesarean section and kept under axenic conditions in Trexler-type plastic isolators for at least 5 generations. The sterility was controlled as previously described (Schwarzer, Makki et al. 2016). Briefly, sterility was assessed every two-weeks by confirming the absence of bacteria, moulds and yeast by aerobic and anaerobic cultivation of mouse feces and swabs from the isolators in VL (Viande-Levure), Sabouraud-dextrose and meat-peptone broth and subsequent plating and aerobic/anaerobic cultivation on blood, Sabouraud and VL agar plates. Conventional (CV) *Helicobacter*-free mice were housed in individually ventilated cages (Tecniplast S.P.A., IT). Experimental animals were obtained by mating of female and male BALB/c mice and their female offspring were weaned at day 28 postnatally. Female offspring were co-housed together until day 60, when they were assigned either to the CV/OVA or CV/Ctrl group, which were housed separately thereafter. Ex-germ-free (exGF) mice were obtained by co-housing of 28-day old GF mice with age-and gender-matched CV mice. GF mice were mono-associated with *Lactobacillus plantarum* WCFS1 (Lp) and the level of bacterial colonization was evaluated weekly by plating serial dilution of feces on de Man, Rogosa and Sharpe (MRS, Oxoid, UK) agar plates as described previously.(Schwarzer, Repa et al. 2011) Colonization remained stable throughout the experiment and reached levels of 2-3 x 10^9^ CFU/g feces. Ceca from control CV, GF, exGF and Lp mice were weighed and a picture was taken. Cecum content was frozen for PCR analysis. Animals were kept in a room with 12 h light-dark cycle at 22°C, fed by OVA-free diet Altromin 1410 sterilized by irradiation and water *ad libitum*. Water was sterilized by autoclaving for GF and Lp-colonized mice. This study was carried out in accordance with the recommendations of the Committee for the Protection and Use of Experimental Animals of the Institute of Microbiology v. v. i., Academy of Sciences of the Czech Republic. All protocols were approved by the same committee.

### Experimental protocol

Female 8-week-old CV, GF, exGF and Lp mice were sensitized *i.p.* within two-week interval with 60 μg of OVA (Worthington, USA) together with 100 μl Alu-Gel-S (Serva, DE) adjuvant and PBS in a final volume of 200 μl on day 1 and 14. Control mice received 100 μl of PBS mixed with 100 μl of Alu-Gel-S. Two weeks after the second *i.p.* sensitization, mice were challenged 8 times at 2–3 day intervals (days 28–44) by *i.g.* gavages of 15 mg OVA in a final volume of 150 μl (OVA groups). Control mice received 150 μl of PBS by *i.g.* gavages (ctrl groups). Diarrhea occurrence was monitored for 30–60 minutes after each *i.g.* exposure. The diarrhea score was assessed according to the following criteria: 0 – normal, well-formed stool, 1 – soft, sticky well-formed stool, 2 – not formed stool, 3 – liquid diarrhea, 4 – more than two episodes of liquid diarrhea after the antigen gavage during the treatment period. The temperature was measured by Thermocouple Thermometer with mouse rectal probe (World Precision Instruments Inc., USA) 30 minutes after the last *i.g.* exposure (**Figure 1A**).

**Figure 1:**
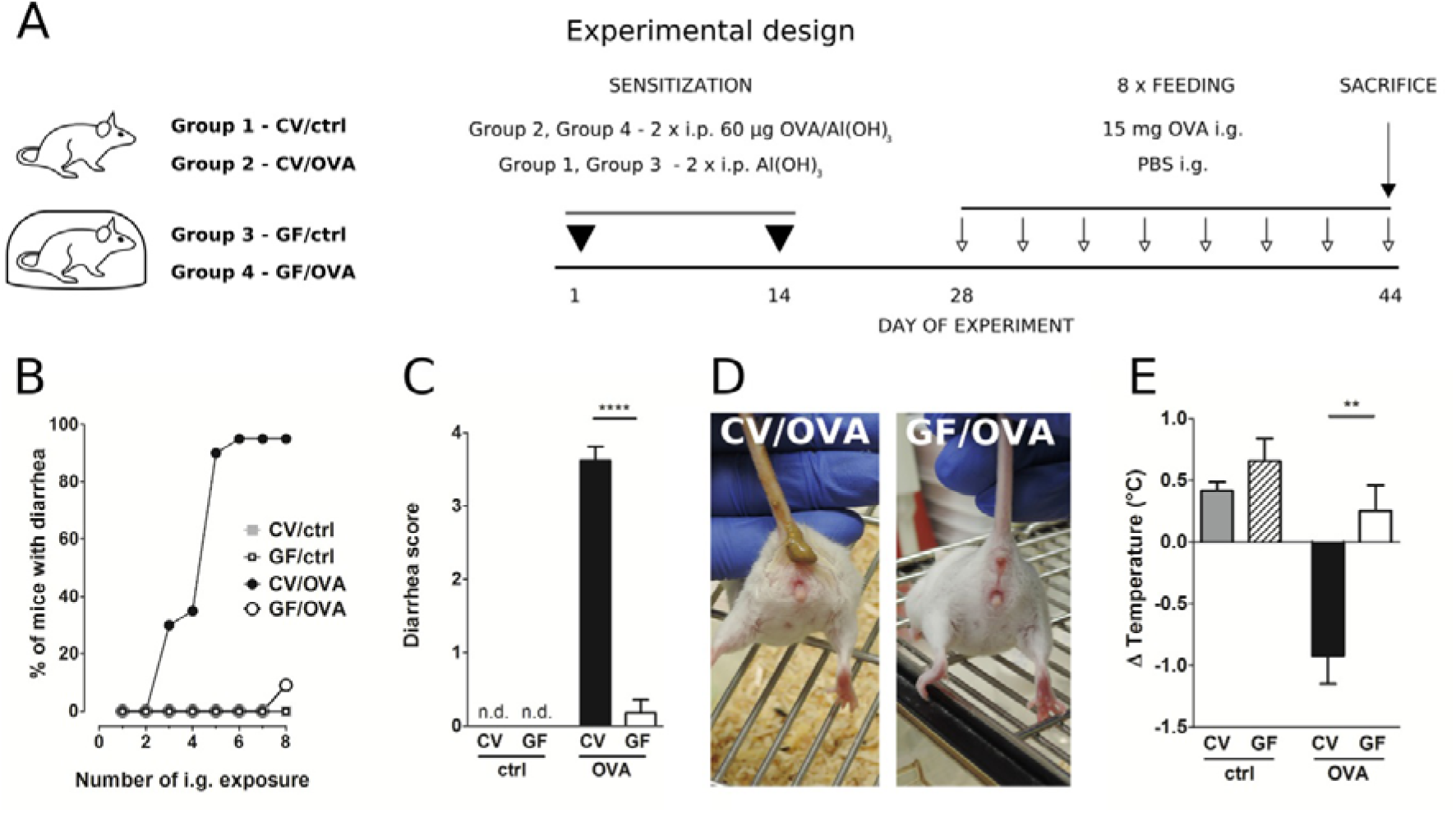
Germ-free mice fail to develop symptoms of OVA-induced food allergy. **A,** Experimental design: Conventional (CV) and germ-free (GF) mice were sensitized twice within a two-week interval by intraperitoneal (i.p.) injection of 60 µg OVA in Al(OH)_3_ (alum) followed by intragastric gavage (i.g.) with 15 mg OVA in PBS 8 times within 17 days (OVA groups). Age-matched control mice were injected with PBS/alum and gavaged by PBS (ctrl groups). After the last i.g. exposure, rectal temperature was assessed and mice were sacrificed by cervical dislocation. Sera and small intestine samples were collected for further analysis. Graphic art was created in freely available professional vector graphics editor Inkscape 0.92 (https://inkscape.org/). **B,** The occurrence of diarrhea was monitored for 60 minutes after each i.g. administration in control CV (grey squares), control GF (open squares), in OVA-treated CV (black circles), and OVA-treated GF (open circles) mice. **C,** The diarrhea score was assessed according to the scoring method described in Material and Methods section. **D,** Representative pictures of CV/OVA and GF/OVA mice after the last i.g. OVA exposure. **E,** The rectal temperature was measured 30 minutes after the last i.g. exposure at day 44 of the experiment in CV/ctrl (grey bars) and GF/ctrl (dashed bars) control mice and in CV/OVA (black bars) and GF/OVA (white bars) OVA-treated mice. Difference in the temperature before and after challenge is shown. Data are plotted as mean values ± SEM. Pooled values of at least two independent experiments (CV/ctrl n = 8, CV/OVA n = 14, GF/ctrl n = 9, GF/OVA n = 11 mice per group) are shown. **P ≤ 0.01, ****P ≤ 0.0001.

### Isolation of bacterial DNA from cecal content and 16S rDNA PCR amplification

Total DNA from 150 mg cecal content was isolated by ZR Fecal DNA kit according to manufacturer‘s instructions (Zymo Research, USA). The purity, integrity and concentration of nucleic acids were confirmed by agarose gel electrophoresis and UV spectrophotometry as previously described. (Srutkova, Spanova et al. 2011) Bacterial 16S rDNA was amplified using PCR with the universal primers 27F (5′ AGA GTT TGA TCC TGG CTC AG 3′) and 1492R (5′ GGT TAC CTT GTT ACG ACT T 3′) as previously described. (Schwarzer, Srutkova et al. 2017) Ten ng of chromosomal DNA from *Escherichia coli* was used as a positive control. Amplification products were separated by electrophoresis in 1.2 % agarose gel, visualized using GelRed^TM^ Nucleic Acid Gel Stain (Biotinum, USA) and images were obtained by Fluorescent Image Analyser FLA-7000 (Fujifilm Corporation, JP).

### Quantification of OVA-specific antibodies

Blood samples were collected at sacrifice and serum was collected after centrifugation. OVA-specific serum IgE, IgG1, IgG2a, and IgA levels were determined by ELISA. (Golias, Schwarzer et al. 2012) Briefly, 96-well microtiter plates were coated with OVA (5 μg/ml). Serum samples were diluted 1/10 for IgE, 1/10000 for IgG1, 1/100 for IgG2, and 1/10 for IgA. Rat anti-mouse IgE, IgG1, IgG2a, and IgA antibodies (1μg/ml, Pharmingen, San Diego, CA, USA) were applied, followed by peroxidase-conjugated anti-rat IgG antibodies (1/1000, Jackson, Immuno Labs, West Grove, PA, USA) for detection. Antibody levels were reported as optical density (OD). The activity of OVA-specific IgE in serum was measured by rat basophil leukemia (RBL) cells degranulation assay as described previously.(Golias, Schwarzer et al. 2012)

### Cellular immune response

At sacrifice, spleens were aseptically removed and single-cell suspensions were prepared in RPMI-1640 containing 10% fetal bovine serum (BioClot GmbH, Aidenbach, Germany) and 1% Antibiotic-Antimycotic solution (Sigma-Aldrich). Cells (6×10^5^/well) were cultured in a flat-bottom 96-well plate (TPP, Trasadingen, Switzerland) without any stimuli or in the presence of OVA (100 mg/well) for 72 hours (37°C, 5% CO2). Supernatants were collected and stored at –40°C until analyses. Levels of IL-4, IL-5, IL-10, IL-13 and IFN-γ were determined by the MILLIPLEX MAP Mouse Cytokine/Chemokine Panel (Millipore, USA) according to manufacturer’s instructions and analyzed with the Bio-Plex System (Bio-Rad Laboratories, USA). Values are reported in pg/ml after subtraction of baseline levels of non-stimulated cell cultures.

### ELISA for mast cell protease-1 and cytokines in jejunal homogenates

Jejunum was aseptically removed and homogenate was prepared as followed. Protease inhibitor (Roche, DE) supplemented with 0.5% Triton X (Sigma-Aldrich, USA) was added to jejunum samples in the ratio 9:1 (w/w). After cooling on ice, the jejunum was homogenized for 1 min/ 40 Hz using Tissue Lyzer and stainless steel beads 7 mm (Qiagen, DE), frozen in liquid nitrogen, thawed and homogenized again. Supernatants were collected after centrifugation and stored at −80°C. Protein content of the homogenates was determined by the Pierce™ BCA Protein Assay Kit (ThermoFisher Scientific, USA) using albumin as a standard. Levels of mouse mast cell protease-1 (MCPT-1) in serum and jejunal homogenates was determined by commercial kit Ready-SET-Go!^®^ (eBioscience, USA) according to manufacturer’s instructions. Levels of IL-4, IL-13 and TNF-α in jejunal homogenates were measured by the MILLIPLEX MAP Mouse Cytokine/Chemokine Panel (Millipore, USA) according to manufacturer’s instructions and analyzed with the Bio-Plex System (Bio-Rad Laboratories, USA). MCPT-1 and cytokine levels in jejunal homogenates are represented per 1 mg of total protein.

### Flow-cytometry analysis

Naive CV and GF BALB/c mice were euthanized by isofluran and peritoneal cavity was washed twice with 5 ml of cold PBS containing 0.1% sodium azide and 0.2% gelatin from cold water fish skin (PBS-gel) (Sigma-Aldrich, USA). Cells (10^6^/well) were blocked for 10 minutes at 4°C in dark by 20% rat heat-inactivated serum in PBS containing 0.1% sodium azide. Staining was performed with fluorochrome labeled anti-mouse monoclonal Abs: FcεRIα-phycoerythrin (BioLegend, USA; clone MAR-1) and CD117(c-kit)-APC-eFluor^®^ 780 (eBioscience, USA; clone 2B8) according to the manufacturer’s recommendation. After 30 min cells were washed four times by cold PBS-gel and data were acquired by FACSCalibur (BD Immunocytometry Systems, Mountain View, CA) flow cytometer. Analysis was performed using FlowJo software (Tree Star, Ashland, OR, USA).

### Cutaneous activation of Mast cells

Intraplantar injection was performed with compound 48/80 (Sigma-Aldrich, USA; 0.5 µg/10 µl per paw) or PBS alone (10 µl per paw) into hind footpad of naive GF and CV mice. Changes in paw width were measured with digital caliper (±0.01 mm; Festa, Czech Republic) at time 0, 30 and 60 minutes. The baseline paw widths (time 0) for each mouse were measured immediately after injection and subtracted from paw widths after application to calculate tissue edema.

### Histology

Intestinal tissue specimens were fixed with 4% paraformaldehyde for 24 hours followed by storage in 80% ethanol. Right ears were fixed in Carnoy’s fluid for 30 minutes and transferred to 96% ethanol. Collected and fixed tissue specimens were dehydrated by using increasing concentrations of ethanol and transferred into methyl salicylate, benzene, benzene-paraffin and paraffin. Sections (5 μm) were deparaffinized in xylene and rehydrated through an ethanol to water gradient and stained for chloracetate esterase activity which is characteristic for mast cell granula. Reagent solution was prepared by mixing of 4% pararosaniline, 2 mol/l HCl, 4% aqueous sodium nitrite, 0.07 mol/l phosphate buffer (pH 6.5), and substrate solution (Naphthol AS-D chloroacetate dissolved in N-dimethylformamide) (all Sigma-Aldrich, USA). The sections were stained by reagent solution for 30 minutes in the dark and counterstained with hematoxylin for 2 minutes. The numbers of mast cells per randomly selected villi in jejunum and per 0.1 mm^2^ in the ear tissue were determined.

### RNA isolation and Real-Time PCR

Jejunal tissues were stored in RNA-later reagent (Sigma-Aldrich, USA) overnight at 4°C and kept at −80°C until processed. Tissue samples were homogenized by Precellys 24 tissue homogenizer (Bertin Technologies, FR) at 5000 rpm for 20 seconds using tubes with zirconium oxide beads. RNA was isolated via RNeasy Mini kit (Qiagen, Valencia, CA). An iScript cDNA Synthesis Kit (BioRad Laboratories, USA) was used to generate cDNA. Real-Time (RT) PCR was performed on the LightCycler^®^ 480 instrument (Roche, DE) using LightCycler^®^ 480 SYBR Green I Master according to the manufacturer’s instructions (Roche, DE). β-actin was used as an internal control to normalize gene expression using the 2-ΔCt method. (Livak and Schmittgen 2001) RT PCR primer sequences are listed in **Supplementary Table 1**.

### Statistical analysis

Statistical analysis between multiple groups was performed by ANOVA with Tukey’s multiple comparison test. Differences between two groups were evaluated using t-test. GraphPad Software was used to evaluate the data (GraphPad Prism 5.04, USA); P values < 0.05 were considered significant. Data are expressed as means ± SEM.

## RESULTS

### Germ-free mice exhibit reduced susceptibility to experimental food allergy

After intraperitoneal injection of OVA adsorbed to alum, CV and GF mice were challenged by oral gavage of OVA eight times over the period of two weeks (**Figure 1A**). Sensitized CV mice were most likely to develop experimental FA after five OVA challenges, with approximately 90% of these animals exhibiting allergic diarrhea (**Figure 1B**). In contrast, none of the GF mice developed diarrhea at this stage. Only after the eighth dose was diarrhea detected in 10% of GF animals (**Figure 1B**). This is reflected in the allergic diarrhea score where the majority of sensitized CV animals exhibited more than two episodes of liquid diarrhea after antigen gavage during the treatment period (**Figures 1C,D**). Sensitized CV mice also exhibited reduced core body temperature after the eighth OVA challenge. This was not observed in the GF animals (**Figure 1E**).

### High levels of Th2-accociated specific serum antibodies and spleen cytokines do not correlate to the low susceptibility of germ-free mice to develop food allergy

Systemic allergic sensitization and oral challenge with OVA led to the induction of -specific IgE, IgG1, IgG2a, and IgA in sera in both experimental groups (**Figures 2 A-E**). In agreement with previous studies (Stefka, Feehley et al. 2014, Kozakova, Schwarzer et al. 2016, Schwarzer, Srutkova et al. 2017), sensitized GF mice exhibited increased levels of OVA-specific IgE in comparison to CV animals (**Figures 2A,B**). The functionality of the OVA-specific IgE was tested in a rat basophil leukemia cell degranulation assay. Sera from sensitized and challenged GF animals induced higher levels of β-hexosaminidase release in comparison to sera from CV mice (**Figure 2B**). While levels of IgG1 were comparable between CV and GF mice (**Figure 2C**), higher specific IgG2a (**Figure 2D**) and IgA (**Figure 2E**) levels were detected in mice raised in the presence of microbiota compared to GF animals. Systemic allergen-specific cellular responses were evaluated by re-stimulating splenocytes with OVA *ex vivo* (**Figures 2F-J**). Stimulation of cells derived from sensitized and challenged GF mice led to the induction of Th2 cytokines (**Figures 2G-J**). Levels of IL-5 and IL-13 were significantly higher in GF mice compared to CV mice (**Figures 2H,I**). A lack of bacterial exposure was associated with reduced levels of OVA-specific IFN-γ production (**Figure 2F**).

**Figure 2:**
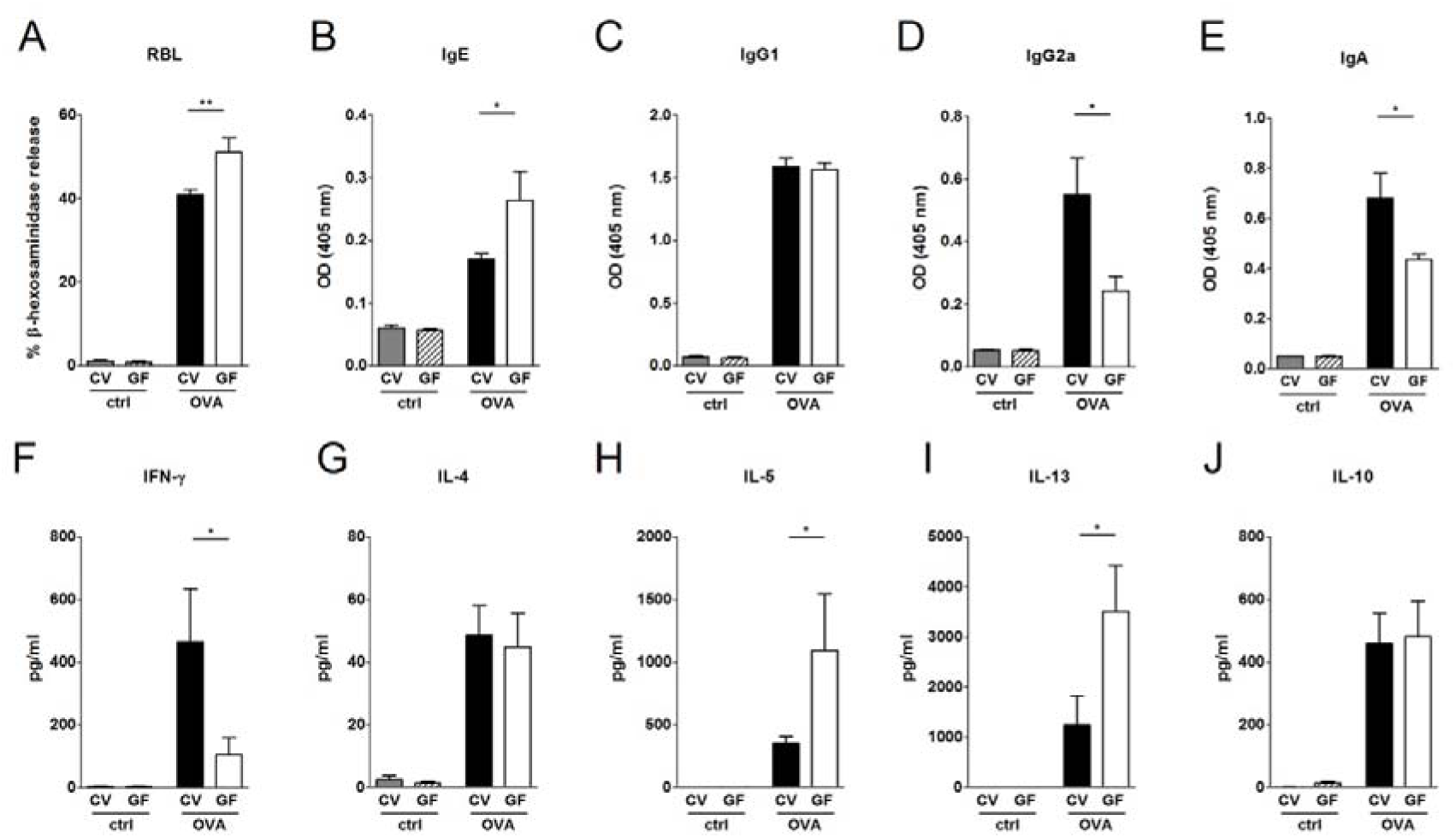
Impact of bacterial colonization on the levels of OVA-specific antibodies in sera and on OVA-induced cytokine production in splenocytes. Levels of OVA-specific antibodies were measured in sera of control CV (grey bars) and GF (dashed bars) mice and sera of OVA-treated CV (black bars) and GF (white bars) mice. **A,** Functional IgE in serum was measured by OVA-mediated β-hexosaminidase release from rat basophil leukemia cells (RBL). OVA-specific **B,** IgE, **C,** IgG1, **D,** IgG2a and **E,** IgA were measured by ELISA and expressed as optical density (OD). Levels of **F,** IFN-γ, **G,** IL-4, **H,** IL-5, **I,** IL-13 and **J,** IL-10 in splenocyte culture supernatants were measured by MILLIPLEX Cytokine panel. Cytokine levels are expressed after subtraction of base line levels of unstimulated splenocytes. Data are plotted as mean values ± SEM. Pooled values of at least two independent experiments (CV/ctrl n = 8, CV/OVA n = 14, GF/ctrl n = 9, GF/OVA n = 11 mice per group) are shown. *P ≤ 0.05, **P ≤ 0.01.

### Absence of microbial colonization leads to a lower density of mast cells in the gut and reduced levels of local and systemic MCPT-1

Given the fact that GF mice were protected from the development of food allergy despite their ability to produce large amounts of OVA-specific IgG1 and IgE as well as pro-allergic systemic cellular responses, we hypothesized that the lack of microbial stimulation resulted in a non-functional effector compartment in the gut. It has been well established that gastrointestinal symptoms during oral antigen-induced anaphylaxis depend not only on IgE but also on the numbers of MC in the intestine (Ahrens, Osterfeld et al. 2012). Here we show that sham-treated GF mice displayed significantly lower numbers of mucosal MC compared to CV mice, as reflected by the lower levels of MCPT-1 in the intestinal tissue (**Figures 3A,B**). This phenomenon, although not as pronounced, was also confirmed in the skin where GF mice harbored 20% fewer MC (**Figures S1A,B in Supplementary Material**). Although the numbers of intestinal MC increased both in CV and GF mice after OVA-treatment, the numbers were significantly lower in GF mice compared to CV mice (**Figures 3A,B**). This was associated with lower expression of intestinal CXCR2 and its ligands CXCL1 and CXCL2, markers associated with the recruitment of MC to the gut (**Figures 3C-E**). Concomitantly, the local production of Th2-associated IL-4 and IL-13 and pro-inflammatory TNF-α was significantly lower in the jejunum of sensitized and challenged GF mice compared to CV animals (**Figures S2A-F in Supplementary Material**). Levels of MCPT-1 in the gut and in serum were significantly lower in GF mice compared to CV mice (**Figures 3F,G**). Although the numbers of intestinal MC in OVA-treated GF mice reached approximately 60% of those detected in CV mice, the production of MCPT-1 in the jejunum reached only 30% of the levels detected in OVA-treated CV mice. These observations suggest that also the maturation status and functionality of MC are influenced by microbiota.

**Figure 3:**
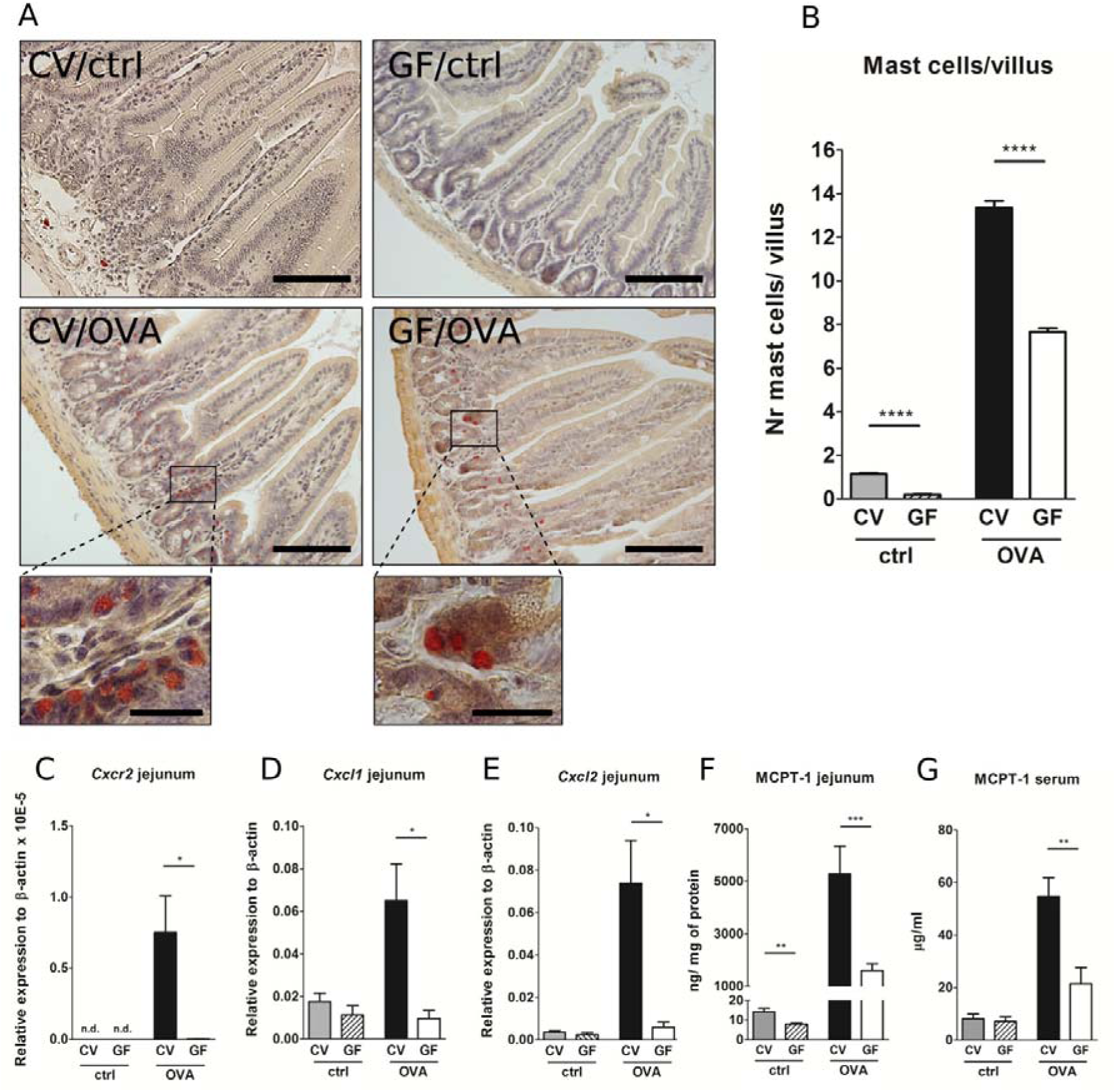
Germ-free mice exhibit low numbers of intestinal mast cells, low expression of *Cxcr2* and its ligands and low levels of Mast cell protease-1 after OVA-sensitization and challenge. **A,** Histological staining of jejunal sections for mastocytosis by hematoxylin/eosin/pararosaniline was performed on samples from control conventional (CV/ctrl) and germ-free (GF/ctrl) and from OVA-treated CV (CV/OVA) and germ-free (GF/OVA) mice (scale bars, 100 μm; inset scale bars, 25 μm) **B,** Quantification of mast cells per villus in jejunal sections (CV/ctrl n = 3, CV/OVA n = 8, GF/ctrl n = 5, GF/OVA n = 5 mice per group). Messenger RNA expression of **C,** *Cxcr2* **D,** *Cxcl1* and **E,** *Cxcl2* (in the jejunal tissues was determined by real-time PCR. Relative expression to β-actin is shown (CV/ctrl n = 6, CV/OVA n = 7, GF/ctrl n = 6, GF/OVA n = 5 mice per group). Mast cell protease-1 (MCPT-1) levels were determined in **F,** jejunal homogenates and in **G,** sera of control conventional (CV; grey bars) and germ-free (GF; dashed bars) mice, and in OVA-treated CV (black bars) and GF (white bars) mice by ELISA. Data are plotted as mean values ± SEM. Pooled values of at least two independent experiments (CV/ctrl n = 8, CV/OVA n = 14, GF/ctrl n = 9, GF/OVA n = 11 mice per group) are shown. **P ≤ 0.01, ***P ≤ 0.001, ****P ≤ 0.0001.

### Lack of microbial colonization impacts the maturation of mast cells

In order to further investigate the role of the microbiota in MC maturation and function, footpads of GF and CV were injected with compound 48/80, which has been used to induce mast cell degranulation and is associated with tissue edema *in vivo* (Chatterjea, Wetzel et al. 2012). The injection of 48/80 induced edema in footpad of CV mice, which was significantly less pronounced in GF animals (**Figure 4A**). Furthermore, the granularity of FcεRIα^+^ and CD117^+^ MC collected by peritoneal lavage was measured by flow cytometry in order to distinguish between immature (SSC-low) and mature (SSC-high) cells. The data show that GF mice have significantly lower numbers of mature MC, and significantly higher numbers of immature MC compared to CV mice (**Figure 4B and Figure S3 in the Supplementary Material**).

**Figure 4:**
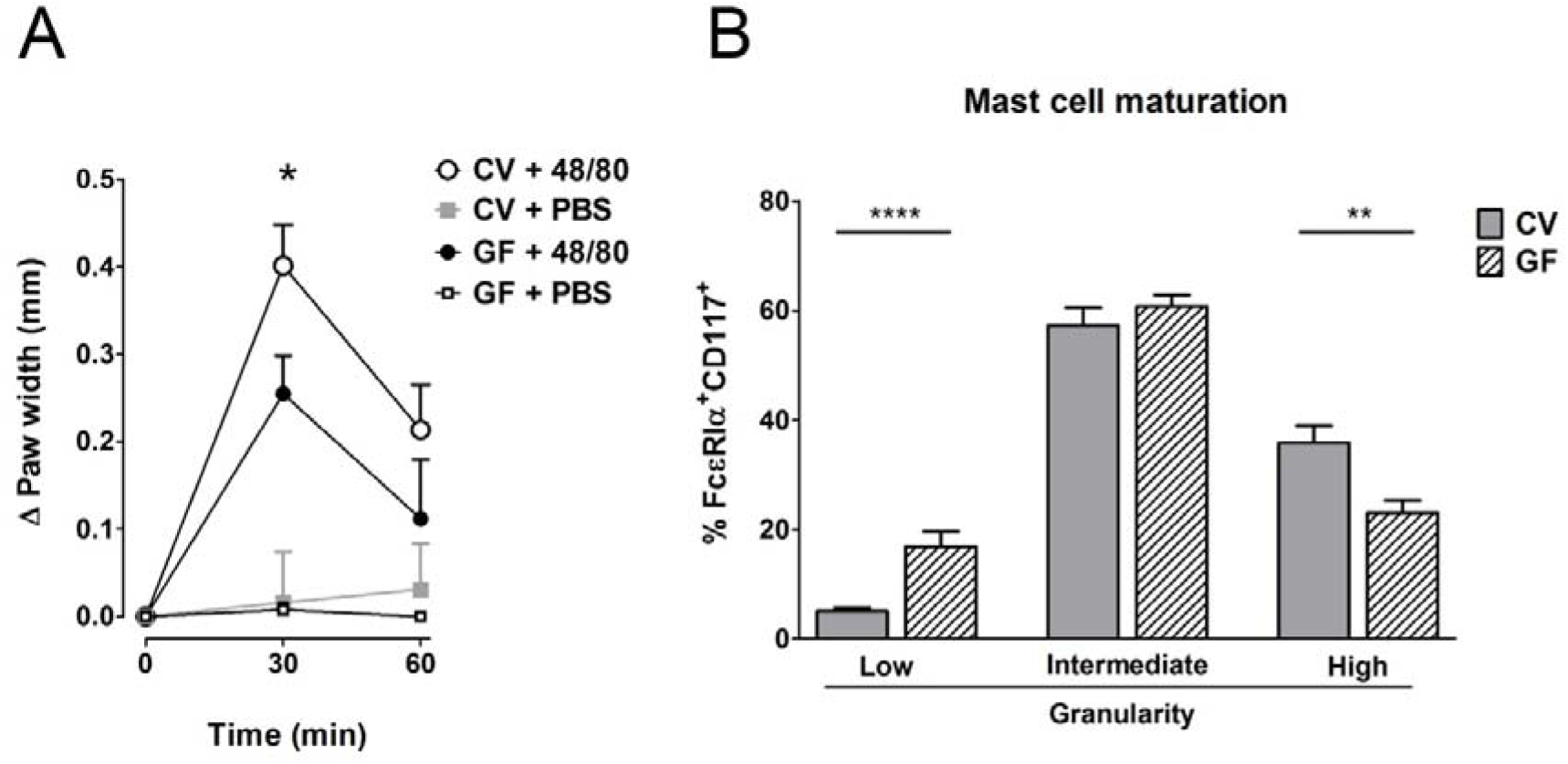
Germ-free mice have less mature mast cells. **A,** GF and CV mice were injected with degranulation inducing compound 48/80 or PBS and edema was recorded. Data are presented as paw width after subtraction of baseline values, GF 48/80 (black circles) n = 6, CV 48/80 (open circles) n = 8, GF PBS (open squares) n = 5, CV PBS (gray squares) n = 8 mice per group. **B,** FcεRIα^+^CD117^+^ polymorphonuclear cells from intraperitoneal lavage were gated and their maturation was assessed according to their granularity. CV (grey bars) n = 9, GF (dashed bars) n = 9 mice per group. Data are plotted as mean values ± SEM. *P ≤ 0.05, **P ≤ 0.01, ****P ≤ 0.0001.

### Reconstitution of germ-free mice with complex microbiota but not with a single commensal strain restores the susceptibility to food allergy

To test whether the susceptibility to food allergy can be restored, we colonized GF mice by co-housing them with CV mice or by gavaging them with human *Lactobacillus* isolate *L. plantarum* WCFS1. Successful conventionalization was verified by decrease in cecal weight and the presence of bacterial DNA measured by PCR in cecal samples (**Figures S4 A,B in the Supplementary Material**). In conventionalized mice (exGF), the incidence of diarrhea, the diarrhea score and the degree of hypothermia reached levels comparable to those observed in CV mice (**Figures 5A-C**). Furthermore, these mice exhibited higher levels of MCPT-1 in the jejunum and serum (**Figures. 5D,E**). In contrast to conventionalized ex-GF mice, GF mice mono-colonized with bacterial strain *L. plantarum* failed to develop clinical symptoms of experimental FA (**Figures 5A-C**). There was no significant difference in the occurrence of diarrhea, diarrhea severity or in the level of hypothermia between Lp mono-colonized mice and GF animals. Moreover, *L. plantarum* did not restore the production of MCPT-1 in the jejunum and serum to those observed in CV mice; levels were similar to those observed in GF mice (**Figures 5D,F**). These data indicate that signals derived from a conventional complex microbiota, rather than a single bacterial strain, are necessary to increase sensitivity to experimental FA.

**Figure 5:**
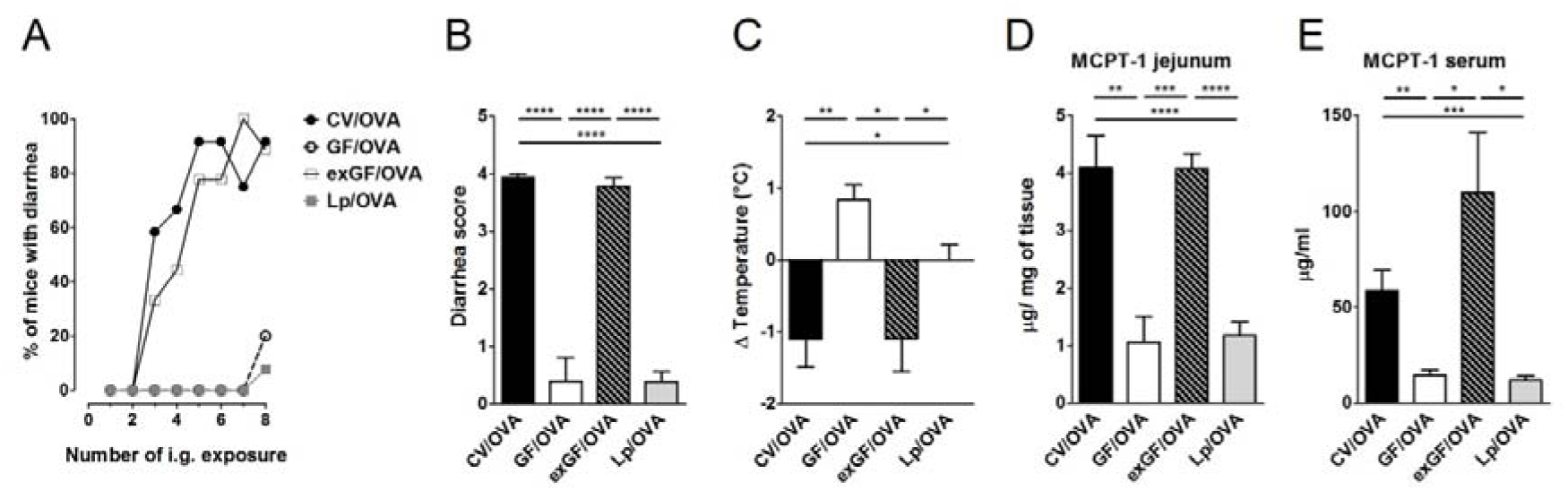
Colonization of germ-free mice by the conventional microbiota but not by single bacterial strain *Lactobacillus plantarum* WCFS1 restores sensitivity to OVA-induced food allergy. Germ-free (GF) mice were colonized by co-housing with age-matched conventional (CV) animals (exGF group) or mono-associated with the strain *L. plantarum* WCFS1 (Lp group). At the age of 8 weeks, all groups were submitted to OVA-sensitization and challenge, as described in Fig.1. **A,** The occurrence of diarrhea during OVA treatment. **B,** The diarrhea score for each group was assessed according to the scoring described in Material and Methods section. **C,** The rectal temperature was taken 30 minutes after the last i.g. exposure at day 44. Difference in the temperature before and after challenge is shown. Mast cell protease-1 (MCPT-1) levels were determined in jejunal homogenates **D,** and in sera **E,** by ELISA. Data are plotted as mean values ± SEM. CV/OVA (n = 8), GF/OVA (n = 5) and pooled values of two independent experiments for exGF/OVA (n = 9) and Lp/OVA (n = 13) colonized mice are shown. *P ≤ 0.05, **P ≤ 0.01, ***P ≤ 0.01, ****P ≤ 0.0001.

## DISCUSSION

Here we report that despite showing high levels of allergic sensitization, GF BALB/c mice are protected against the development of OVA-induced FA, exhibiting low incidence of diarrhea and hypothermia. In the intestine, GF mice displayed reduced numbers of MC and lower levels of MCPT-1 after allergic challenge in comparison to CV mice. Further we confirmed the altered MC maturation and function in the absence of microbiota. Finally, colonization of GF mice with conventional microbiota, but not their mono-association with single bacterial strain *L. plantarum* WCFS1, induced the susceptibility of exGF mice to OVA-induced FA. In accordance with the hygiene hypothesis, GF animals have been shown to develop increased levels of serum IgE after allergic sensitization in comparison to their CV counterparts (Morin, Fischer et al. 2012, Schwarzer, Srutkova et al. 2013, Stefka, Feehley et al. 2014, Kozakova, Schwarzer et al. 2016). Here we show that sensitization and challenge lead to increased OVA-specific IgE levels in sera and increased production of OVA-specific IL-5 and IL-13 in re-stimulated splenocytes from GF mice compared to CV animals. The concept that signals derived from commensal bacteria are crucial to normalize elevated Th2 responses have been tested previously by us and others. For example, colonization of GF mice with conventional or SPF microbiota, with a well-defined mixture of different strains, or with only a single bacterial strain led to reduced levels of serum IgE and Th2 cytokine production in comparison to GF animals (Hill, Siracusa et al. 2012, Cahenzli, Köller et al. 2013, Schwarzer, Srutkova et al. 2013, Kozakova, Schwarzer et al. 2016). Recently, we have expanded this observation to residual bacterial fragments and showed that the presence of LPS in sterile food was sufficient to reduce high levels of allergen-specific IgE in GF sensitized mice (Schwarzer, Srutkova et al. 2017).

Surprisingly, despite the elevated systemic OVA-specific humoral and cellular Th2 responses, GF mice were protected from FA symptoms, i.e. they failed to develop allergic diarrhea and hypothermia. This observation is in disagreement with studies by Cahenzli *et al*, Rodriguez *et al.* and Stefka *et al*. who showed that GF mice are more susceptible to FA (Rodriguez, Prioult et al. 2011, Cahenzli, Köller et al. 2013, Stefka, Feehley et al. 2014). There are several possible explanations for this discrepancy. First, differences in the sensitization and challenge protocols. We used 2 intraperitoneal doses of 60 µg OVA/Alum followed by eight challenges of 15 mg OVA by gavage, as previously established by Golias *et al*. (Golias, Schwarzer et al. 2012). In contrast, Cahenzli *et al*. sensitized mice with one subcutaneous dose of 50 µg OVA/Alum followed by single oral challenge with 50 mg OVA (Cahenzli, Köller et al. 2013). A very different model was used by Stefka *at al*. and Rodriguez *et al*., where mice were sensitized by intragastric gavages with antigen admixed with cholera toxin (CT) (Rodriguez, Prioult et al. 2011, Stefka, Feehley et al. 2014). CT is known as a potent mucosal adjuvant which induces mobilization and maturation of intestinal immune cells (Anjuere, Luci et al. 2004) as well as the induction of Th1/Th2/Th17 responses (Mattsson, Schon et al. 2015). Thus, the application of CT to GF mice might provide signals for maturation or recruitment of MC to the gut tissue. Second, there are differences in the readouts and reporting of clinical symptoms. In Cahenzli *et al*. and in Stefka *et al*., the allergic sensitization and oral challenge did not induce hypothermia in SPF or CV mice (Cahenzli, Köller et al. 2013, Stefka, Feehley et al. 2014). This is surprising since we and others have shown that immunization in the presence of adjuvant followed by oral challenge with or without adjuvant results in clinical symptoms, such as hypothermia or diarrhea in both conventional animals or animals with SPF microbiota (Osterfeld, Ahrens et al. 2010, Ahrens, Osterfeld et al. 2012, Yamaki and Yoshino 2012, Chen, Lee et al. 2015, Lee, Chen et al. 2016). Third, different sterilization methods of chow for GF animals. In our study, GF animals were fed by γ-irradiated chow, which is in contrast to sterilization method used by Cahenzli *et al*., where the chow was autoclaved (Cahenzli, Köller et al. 2013). Autoclaving leads to formation of advanced glycation end-products as a result of chemical reaction between reducing sugars and amino acids in proteins or lipides (Meillard reaction) (Gupta, Gupta et al. 2016). These products has been recently pointed at as one of the possible causes of increasing FA incidence in Western countries (Smith 2017). This is of special interest, as autoclaved, but not irradiated food has been shown to increase the numbers of intestinal MC in GF rats (Meslin, Wal et al. 1990).

To determine the discrepancy between high levels of sensitization and the absence of FA symptoms, we investigated the presence of intestinal MC, which are widely recognized key effectors of allergy in the periphery (Brandt, Strait et al. 2003, Yamaki and Yoshino 2012, Chen, Lee et al. 2015). Here we demonstrate that GF animals exhibit low levels of MC in the gut under homeostatic conditions and that MC numbers are still reduced after allergen sensitization and oral challenge in comparison to CV mice. The reduced numbers of intestinal MC in GF mice were accompanied by decreased expression of *Cxcr2* and its ligands in intestinal tissue. Concomitantly, challenged GF mice had significantly lower production of MCPT-1 both locally in the intestinal tissue and systemically in sera. Further experiments are required to dissect whether the microbiota impacts MC recruitment and/or function directly or indicrectly via the action on intestinal epithelial or innate lymphoid cells.

Together with a previous report about the crucial role of MC for the development of FA symptoms (Brandt, Strait et al. 2003), our data suggests that the MC homing to the intestinal effector compartment is impaired in GF animals. Chen *et al*. have also shown that the severity of FA correlates with intestinal MC numbers as mouse strains without intestinal MC (i.g. C57BL/6 or C3H/HeJ) did not exhibit clinical symptoms of experimental FA after antigen sensitization and challenge (Chen, Lee et al. 2015). However, it has been shown that long-term breeding of different mouse strains in isolation from each other results in distinct and distinguishable intestinal microbial communities (Ubeda, Lipuma et al. 2012). Due to the fact that the different mouse strains used in the study by Chen *et al*. were not littermate controlled, the role of microbiota in the susceptibility to food allergy cannot be ruled out (Chen, Lee et al. 2015).

In our model, intestinal MC numbers in sensitized and challenged GF animals reached 60% of those observed in CV animals. However the GF mice were still fully protected from the development of allergic diarrhea and hypothermia. This indicates that not only the numbers of local MC in GF mice, but also the functionality and maturation status of MC are impaired in these mice. In a recent paper, Wang *et al*. clearly demonstrated that the microbiota drives the recruitment and functionality of skin MC (Wang, Mascarenhas et al. 2016). In order to test the functionality of MC we challenged the GF and CV mice by injection of degranulation compound 48/80. GF mice exhibited significantly lower swelling compared to CV animals confirming the impaired functionality of MC in the absence of microbiota. Next, we addressed the impact of the microbiota on the MC maturation status. Previously Dahlin *et al*. showed that the low granularity of MC corresponds with their low maturation status (Dahlin and Hallgren 2015). We found that there was an increased percentage of MC with low granularity in GF animals compared to CV animals and that these immature MC expressed lower levels of CD117 (data not shown), a known surface maturation marker (Wang, Mascarenhas et al. 2016).

Finally, we could show that conventionalization rendered exGF mice sensitive to FA, as demonstrated by hypothermia, diarrhea, and elevated levels of MCPT-1 in the gut and serum. Interestingly, mice mono-colonized with Gram-positive strain *L. plantarum* remained unresponsive to OVA challenge. This observation is surprising, as *L. plantarum* WCFS1 is a bacterial strain with strong immunomodulatory properties (Rigaux, Daniel et al. 2009, Smelt, de Haan et al. 2012, Gorska, Schwarzer et al. 2014) and oral application of this strain has been shown to aggravate the severity of peanut FA in a mouse model (Meijerink, Wells et al. 2012). Nevertheless it has been well documented that different bacterial strains, and even the strains of the same species, may differ in their immunomodulatory potential (Schabussova, Hufnagl et al. 2011, Srutkova, Schwarzer et al. 2015). Whether mono-colonization by other bacterial strains (e.g. Gram-negative) or supplementing the GF mice with specific bacterial products (e.g. LPS, peptidoglycan, lipoteichoic acid) can impact the maturation and function of intestinal MC and render the mice susceptible to food allergy remains to be determined. Taken together, we report here that commensal bacteria impact MC migration and maturation, thus playing a key role in the susceptibility to food-induced allergy. Our model based on CT-free systemic sensitization and oral challenge leads to full development of hypothermia and diarrhea in CV settings but not GF mice, represents an important tool to investigate the role of microbiota in the development or prevention of infectious or immune-mediated inflammatory diseases. The mechanistic insight into the role of the commensals-MC-FA axis, with a focus on the microbiota-induced recruitment and maturation of MC in the intestinal mucosa, can pave the way to the design of novel strategies for the prevention and treatment of food allergy in humans.

## ACKNOWLEDGMENTS

Excellent technical assistance of J. Jarkovska, A. Smolova, I. Grimova, B. Drabonova, D. Drasnarova and K. Ambroz is gratefully acknowledged. Authors would like to thank Dr. Constance Finney for critical reading and edition of the manuscript.

## AUTHORS CONTRIBUTIONS

M.S., L.T., H.K, and I.S. conceived and designed the experiments. M.S., P.H., J.G., T.H., D.S., J.A. and CZ performed the experiments. M.S., P.H., J.G., T.H., D.S. analyzed the data. M.S., P.H., U.W., L.T., H.K., and I.S. wrote the paper. All authors reviewed the manuscript.

## Supplementary Material

**Table S1.**
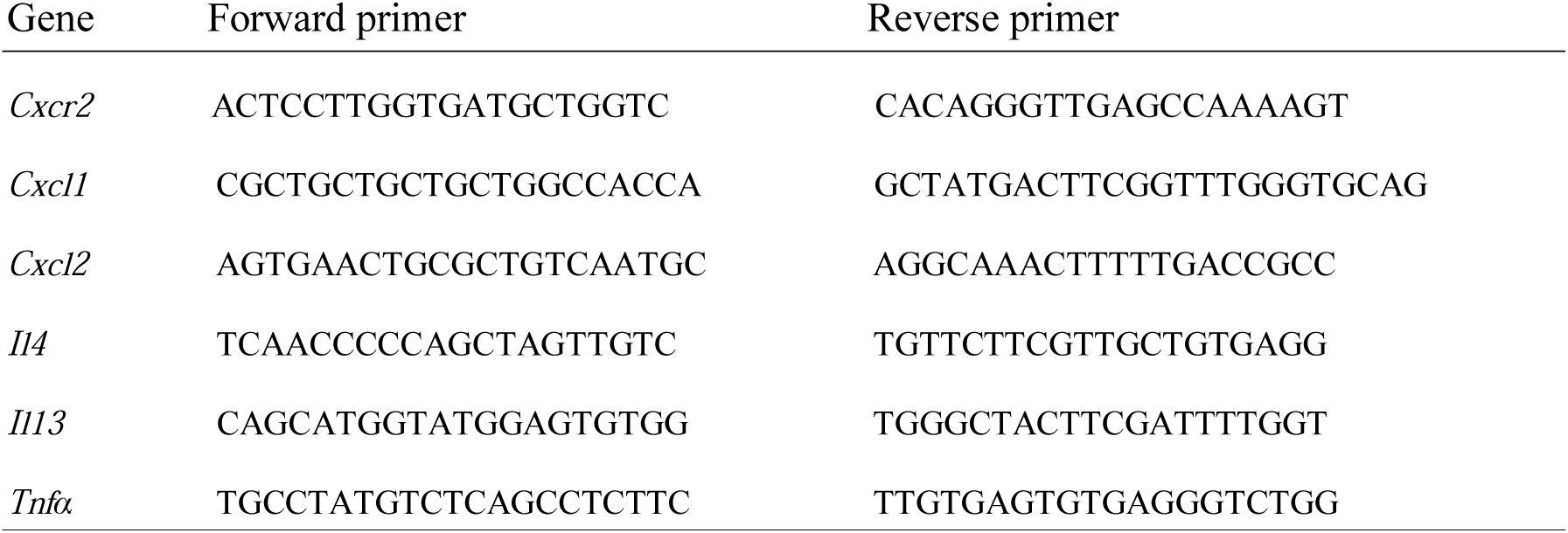
Sequences of real-time PCR primer.

**Figure S1.**
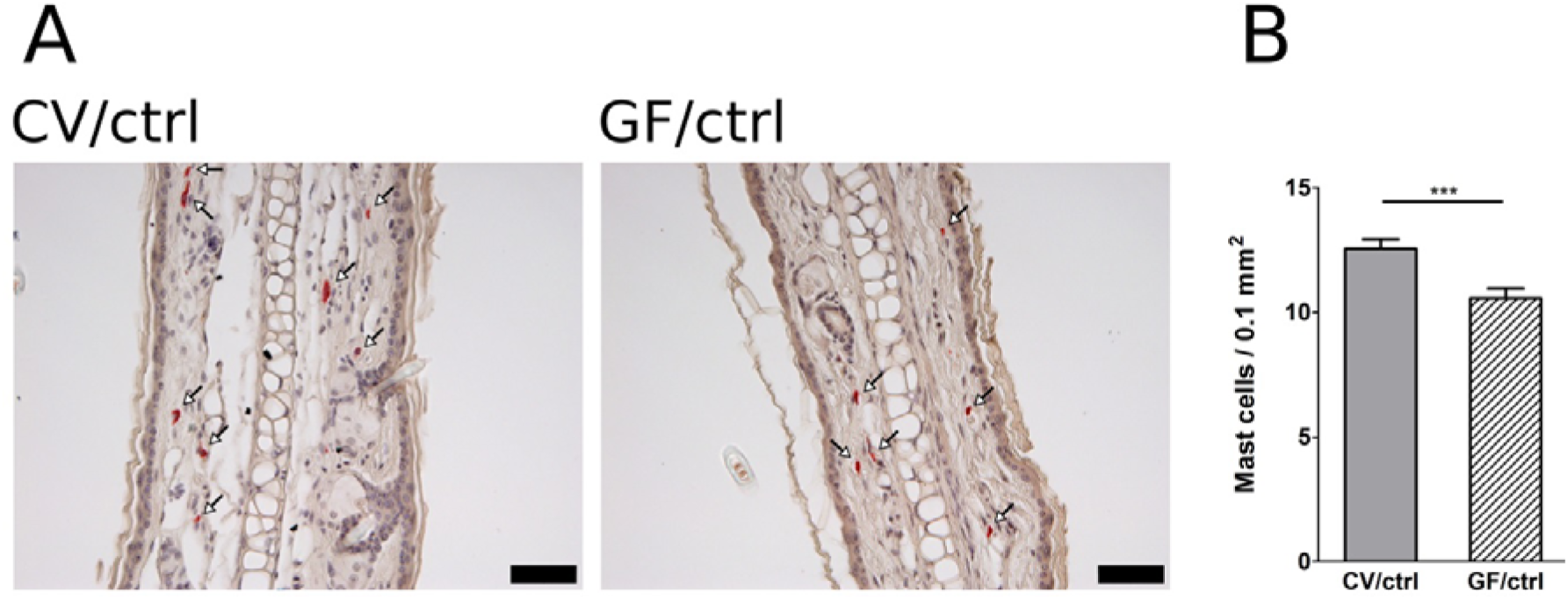
Germ-free mice exhibit reduced numbers of skin mast cells compared to conventional mice. **A,** Histological staining of ear sections for mastocytosis by hematoxylin/eosin/pararosaniline was performed on samples from control conventional (CV/ctrl) and germ-free (GF/ctrl) animals, (scale bars, 50 μm). **B,** Mast cells were quantified per 0.1 mm2 area of the tissue. Pooled values of n = 5 mice per group are shown. The occurrence of mast cells in 8-12 independent areas were counted for each individual mouse.***P ≤ 0.001.

**Figure S2.**
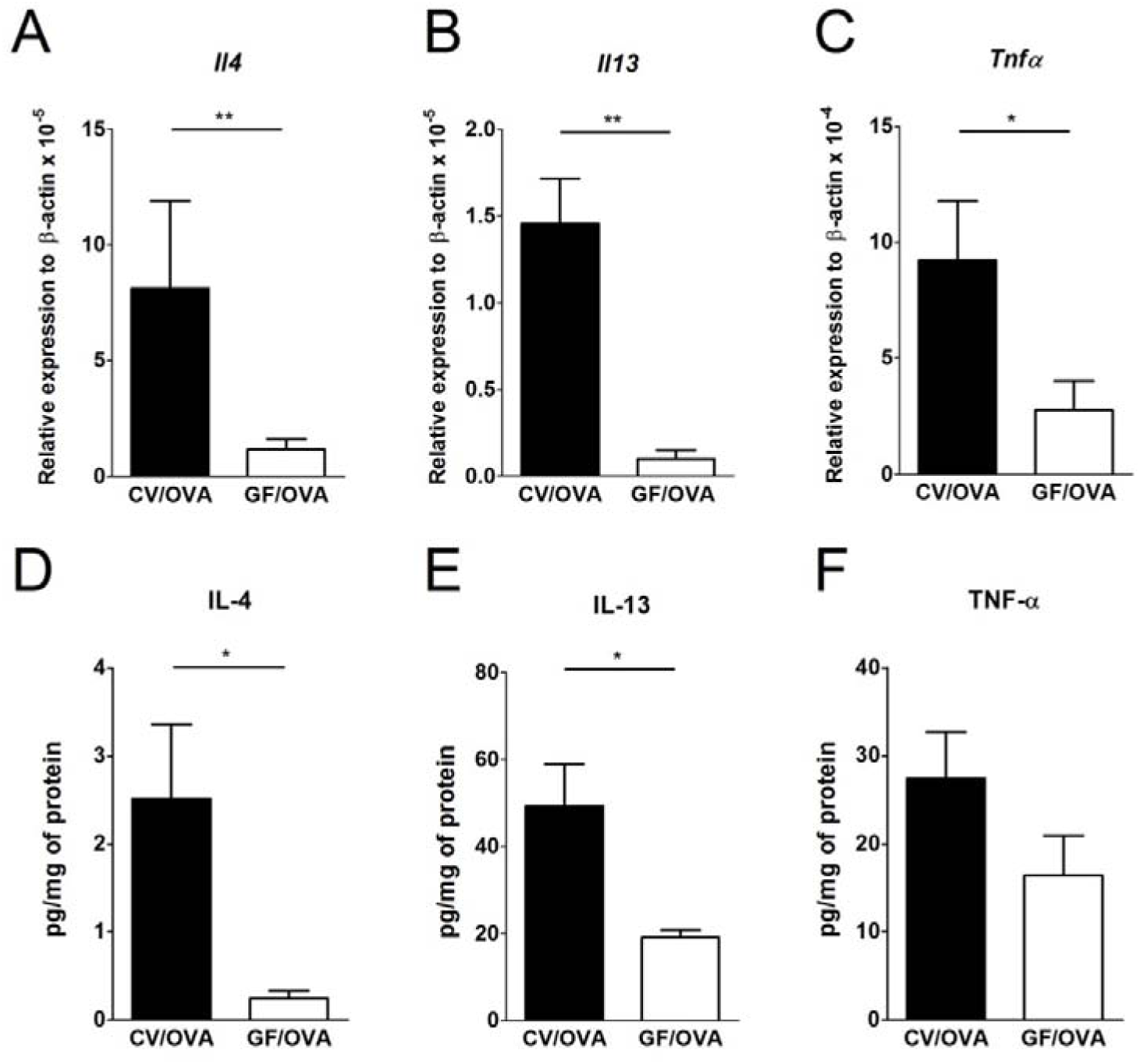
Messenger RNA and protein levels of IL-4, IL-13 and TNF-α are low in the jejunum of OVA-sensitized and challenged germ-free mice. **A,** Messenger RNA expression of *Il4*, **B,** *Il13* and **C,** *Tnf*α (in the jejunal tissues of OVA-treated conventional (CV/OVA; black bars; n = 7) and germ-free (GF/OVA; white bars; n = 5) mice was determined by Real-Time PCR. Relative expression to β-actin is shown. Protein levels of IL-4 (**d**) IL-13 (**e**) and TNF-α (**f**) in the jejunal tissue homogenates of OVA-treated CV (black bars, n = 14) and GF (white bars, n = 11) mice were determined by ELISA. Levels are normalized per 1 mg of protein. Data are plotted as mean values ± SEM. *P ≤ 0.05, **P ≤ 0.01.

**Figure S3.**
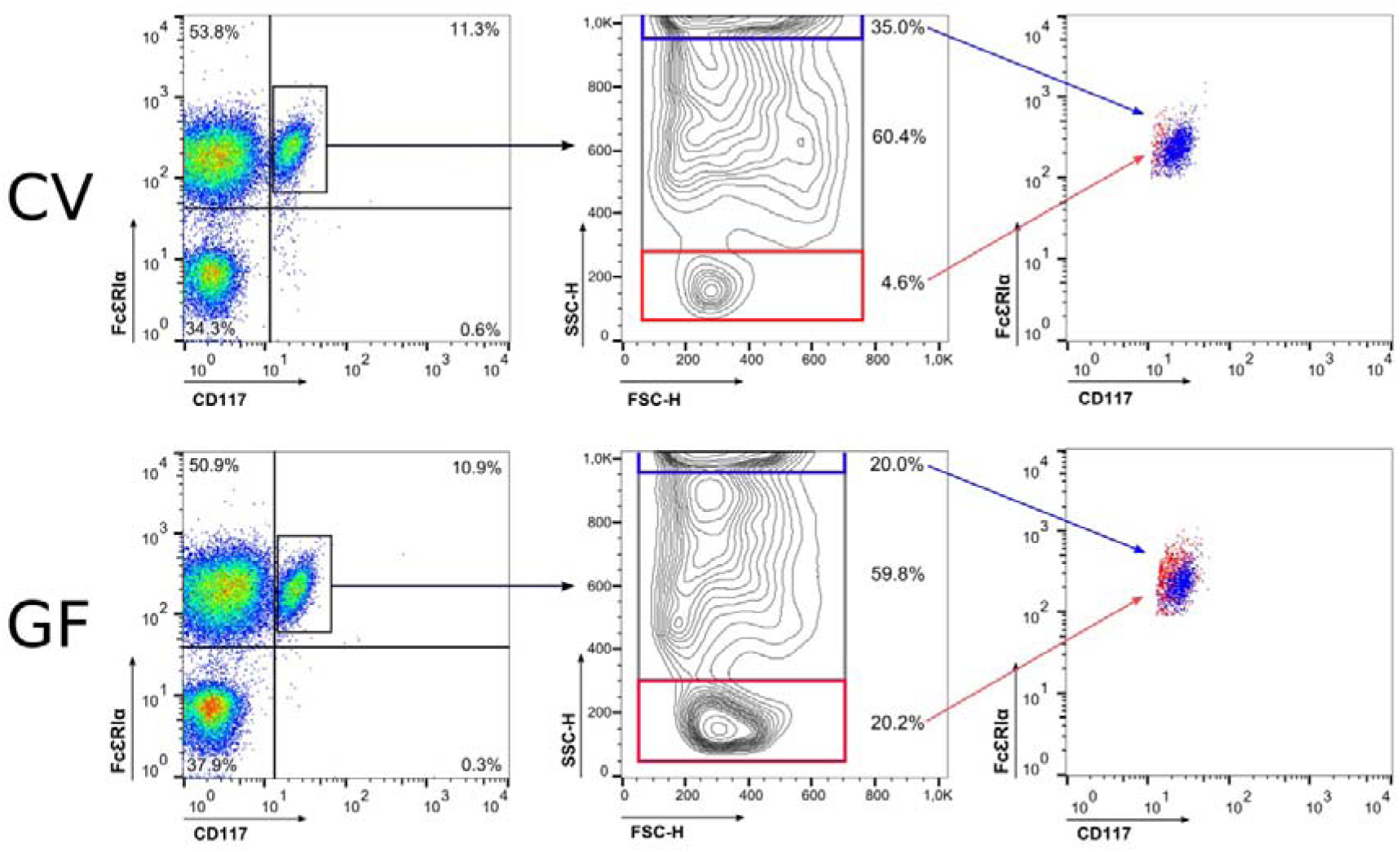
Flow cytometry analysis of peritoneal mast cells. **A,** Cells from intraperitoneal lavages from CV (n = 9) and GF (n = 9) mice were analyzed. Size (forward scatter - FSC) and granularity (side scatter - SSC) of FcεRIα and CD117 (ckit) positive cells (mast cells) out of total polymorphonuclear leucocytes were analyzed and displayed using counter plot. Three populations (low, intermediate and high) were established according to cell granularity and the percentage of their density was analyzed. **B,** Median fluorescence intensity (MFI) of the CD117 expression are shown for SSC-high (blue dots) and SSC-low (red dots) mast cell populations from CV (n = 9) and GF (n = 9) mice. Horizontal bar represents mean values ± SEM. ***P ≤ 0.001, ****P ≤ 0.0001.

**Figure S4:**
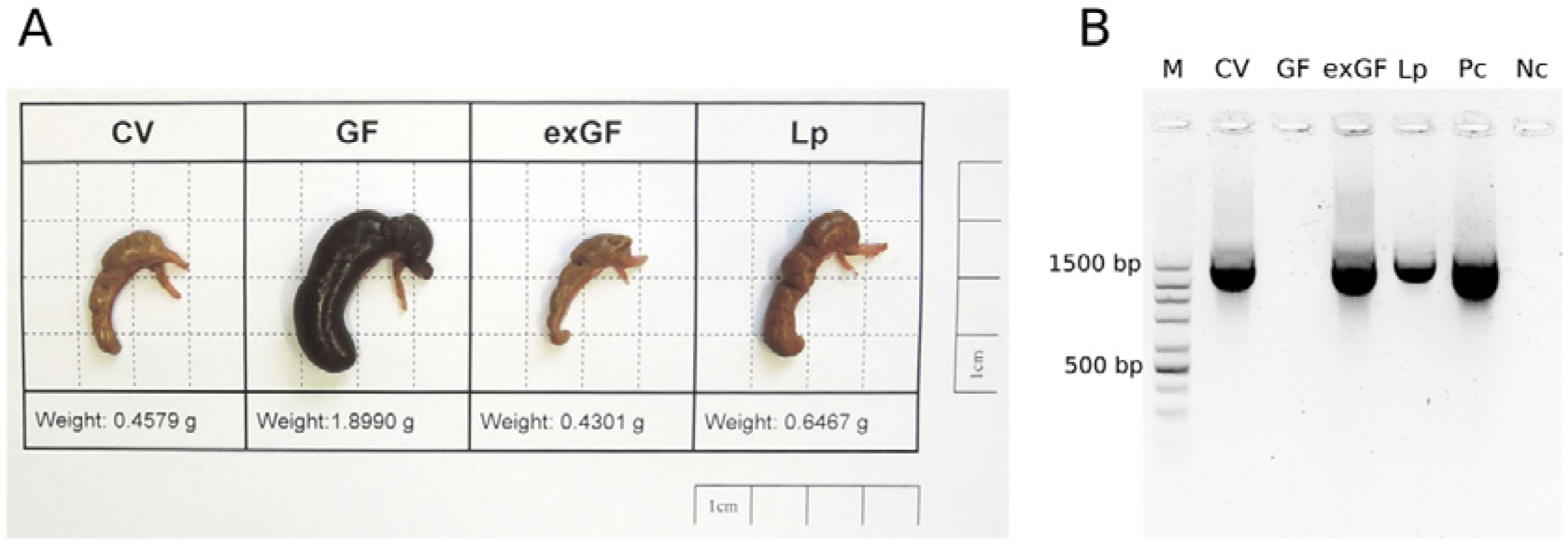
Successful colonization of germ-free mice with the conventional microbiota and with the *Lactobacillus plantarum* WCFS1. **A,** Representative ceca from conventional (CV), germ-free (GF), exGF (GF mice colonized by cohousing with age-matched CV animals) and *L. plantarum* WCFS1-monocolonized gnotobiotic (Lp) mice were dissected at the end of experiment, weighed and photographed. **B,** Total DNA from cecal content was isolated and PCR was performed with bacteria-specific primers. PCR products were separated by electrophoresis in 1.2% agarose gel. 10 ng of *E. coli* DNA was used as positive control (Pc), H2O as negative control (Nc), marker (M).

## Abbreviations used

CV: Conventional
ExGF: GF mice co-housed with conventional mice
FA: Food allergy
GF: Germ-free
IL: Interleukin
LPS: Lipopolysaccharide
MC: Mast cells
MCPT-1: Mast cell protease-1
OVA: Ovalbumin
PCR: Polymerase chain reaction
SPF: Specific pathogen free

